# Unsupervised determination of lung tumor margin with widefield polarimetric second-harmonic generation microscopy

**DOI:** 10.1101/2022.07.25.501438

**Authors:** Kamdin Mirsanaye, Leonardo Uribe Castaño, Yasmeen Kamaliddin, Ahmad Golaraei, Lukas Kontenis, Edvardas Žurauskas, Roya Navab, Kazuhiro Yasufuku, Ming-Sound Tsao, Brian C. Wilson, Virginijus Barzda

## Abstract

The extracellular matrix (ECM) is amongst many tissue components affected by cancer, however, morphological changes of the ECM are not well-understood and thus, often omitted from diagnostic considerations. Polarimetric second-harmonic generation (P-SHG) microscopy allows for visualization and characterization of collagen ultrastructure in the ECM, aiding in better understanding of the changes induced by cancer throughout the tissue. In this paper, a large region of hematoxylin and eosin-stained human lung section, encompassing a tumor margin, connecting a significant tumor portion to normal tissue was imaged with P-SHG microscopy. The resulting polarimetric parameters were utilized in principal components analysis and unsupervised k-means clustering to separate normal- and tumor-like tissue. Consequently, a pseudo-color map of the clustered tissue regions is generated to highlight the irregularity of the ECM collagen structure throughout the region of interest and to identify the tumor margin.

## Introduction

Lung cancer is the deadliest form of cancer, which was responsible for 18% of worldwide cancer-related deaths in 2020 [1]. Currently, histological diagnosis of cancer is based on examination of hematoxylin and eosin-stained (H&E) tissue sections under a white light microscope. The tumor and its properties are determined by immunohistochemical and molecular examination of the tumor cells. However, the tumor microenvironment has not been evaluated in detail at present, and its effect on the biological behavior of the tumor is poorly understood. The structure and organization of collagen fibers within the extracellular matrix (ECM) have shown to become significantly modified due to the presence of tumor in tissue. Although several tissue stains have been developed and tested to study collagen in histological tissues [3, 4, 6, 5], collagen morphological characteristics have not gained widespread use as a potential cancer biomarker. Polarimetric Second-Harmonic Generation (P-SHG) microscopy is a second order nonlinear optical imaging technique that enables visualization and structural characterization of noncentrosymmetrically arranged molecules, such as collagen. P-SHG was extensively used in laser scanning microscopy systems to differentiate between normal and diseased tissues, including various types of malignancies in lung [10], breast [12, 13, 14], ovary, pancreas [15], and thyroid [11]. This microscopy technique provides detailed ultrastructural information for every voxel of the imaged biological sample in the form of nonlinear susceptibility tensor element ratios. Although powerful in differentiating normal and pathological tissue, laser scanning P-SHG is relatively slow and therefore ill-suited for imaging of large area tissue sections. In a previous study, a widefield P-SHG microscope was developed and utilized for a whole tissue microarray imaging with a fast scan-less polarimetric SHG imaging of individual breast tissue cores of 600*μm* diameter. In tandem with supervised machine-learning, the classification of normal and tumor breast tissue cores was achieved with over 94% accuracy [27]. Consequently, large area and high-throughput P-SHG histopathology investigations may now be performed in a timely manner. In this work, the widefield P-SHG microscope was used together with unsupervised machine learning to map the ECM morphological features across a large section of a non-small cell lung carcinoma tumor histological slide. The technique generated a series of polarimetric and texture parameters that enabled identification of the collagen ultrastructure and organization across the tumor margin. The importance of the extracted parameters were examined with Principal Components Analysis (PCA) and the results were utilized to optimize a k-means clustering algorithm and identify normal- and tumor-like tissue regions. Widefield P-SHG microscopy imaging of large area tissue sections accurately delineated tumor margins based on changes in the ECM collagen in the absence of supervision, using H&E-stained tissue sections which were prepared for the gold-standard histopathologic cancer diagnostics.

## Results

### Large-area P-SHG imaging of lung tissue

The recently developed widefield P-SHG microscope [27] was used to image a large area of lung tissue, encompassing a non-small cell lung carcinoma tumor portion on the left, and extending to normal tissue on the right as shown in the H&E image (Fig. 1a). According to Stokes-Mueller polarimetry described previously for widefield P-SHG [27], 5 distinct polarimetric parameters were computed and used to characterize the ECM. These polarimetric parameters include SHG intensity of all circular incident polarizations, R-Ratio which is comprised of two second-order nonlinear optical susceptibility elements (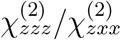, where the z-axis is parallel to the collagen fiber axis), degree of ciruclar polarization (DCP), as well as, SHG circular and linear dichroism (SHG-CD and SHG-LD, respectively).

**Figure 1:**
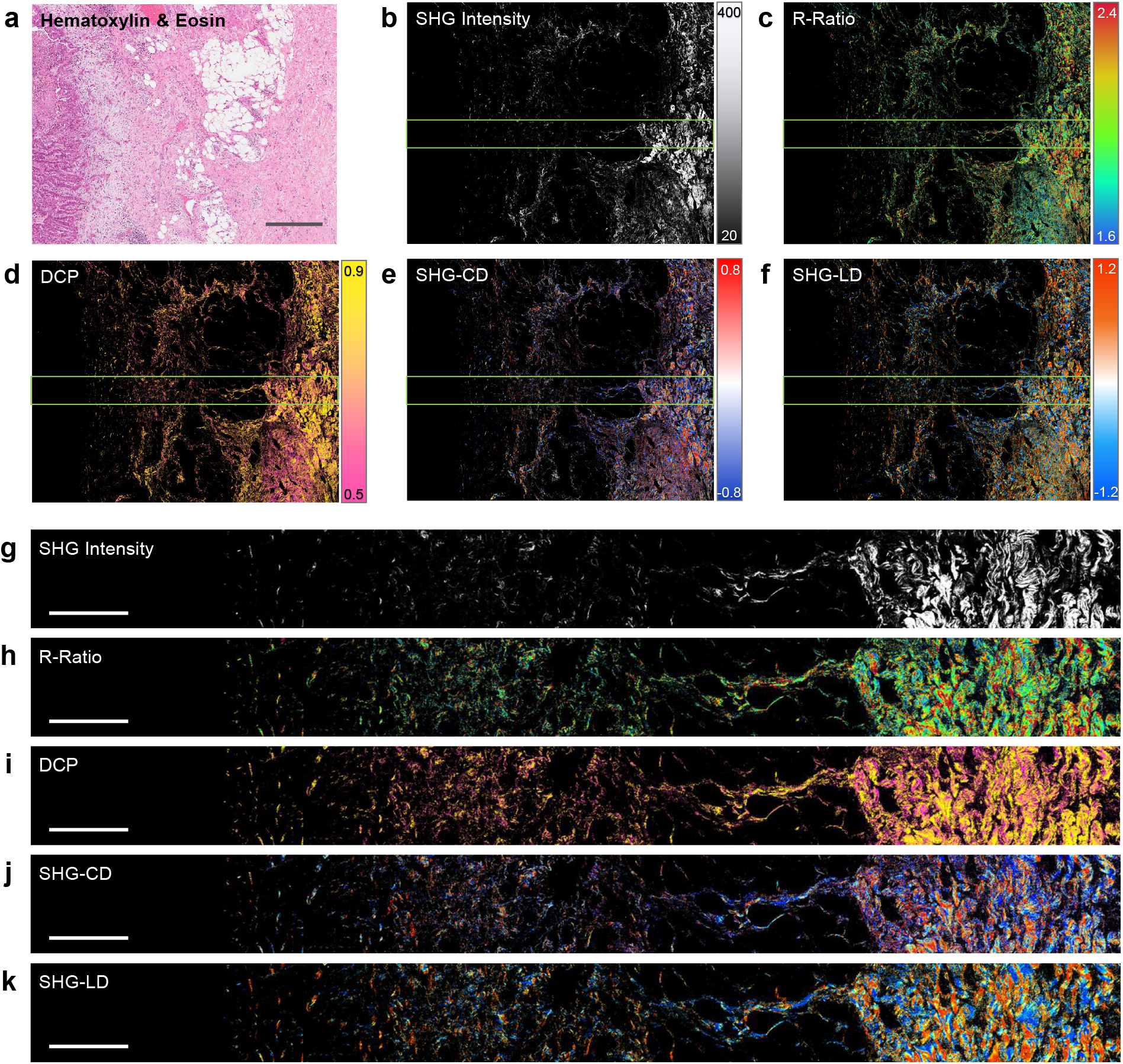
P-SHG microscopy of lung tissue tumor margin. A large region of human lung tissue is depicted in Hematoxilin and Eosin (H&E) stained white-field image (a), Second-Harmonic Generation (SHG) intensity (b), R-ratio (c), Degree of Circular Polarization (d), as well as, SHG circular (e) and linear dichroism (f), respectively. Scale bar: 500*μ*m. Close up regions of the tissue corresponding to area highlighted by green border showcase the variation of polarimetric parameters from tumor (left side) to normal-like (right side) tissue (g-k). Scale bar: 200*μ*m.

The second-harmonic generation efficiency and signal intensity is highly dependent on the organization of polar collagen fibers in the ECM. It was shown that while parallel arrangements of collagen fibers results in a strong SHG signal, anti-parallel arrangements or disordered configurations often significantly reduce the measured SHG intensity [21]. In addition, it was previously illustrated that SHG intensity was significantly reduced in solid tumors, suggesting degradation and structural disorder of collagen fibers in the ECM of tumor tissue [22, 23]. R-Ratio was previously utilized to differentiate between various tissue types, including normal and malignant breast, lung, pancreas, and thyroid tissues [12, 10, 15, 11]. It is important to note that R-ratio varies with molecular composition, as well as orientation of collagen fibers in the ECM. DCP was identified to be proportional to R-ratio for collagen, which suggested depolarization of circularly-polarized SHG signal also varied with collagen fiber molecular composition and orientation. Although mathematically complex, both SHG-CD and SHG-LD are highly dependent on collagen fiber out-of-plane and in-plane orientation, respectively. SHG-CD was shown to be important in highlighting collagen fiber polarity [24, 25, 9], used to study ovarian and breast cancer [26, 27]. Consideration of SHG-LD was recently introduced for P-SHG investigation of breast cancer, showcasing in-plane fiber orientation over a large section of the tissue [27]. Representative large-area pseudo-color maps of all computed polarimetric parameters are shown in Figs. 1b-f, depicting the general observed trends in polarimetric parameter values between normal and tumor tissue. Each large area image is comprised of a 3 × 4 grid of widefield P-SHG images, each containing 2048 × 2048 pixels and spanning a 670*μm* × 670*μm* area (total image size of 2mm × 2.7mm, containing over 50 million pixels). Pixels below signal to noise ratio of 1 were discarded from further analysis and appear black in the images. A narrow band of the polarimetric parameter map, corresponding to the highlighted regions in Figs. 1b-f, are shown in Figs. 1g-k, to better visualize the polarimetric parameter variations from left to right (tumor to normal).

As expected, it is apparent that SHG intensity decreases significantly toward the tumor side of the area of interest. Moreover, the normal side of the tissue section contains a larger number of bright pixels (we will refer to as pixel density). The remaining polarimetric parameters such as R-Ratio, degree of circular polarization (DCP), as well as SHG circular and linear dichroism (SHG-CD and SHG-LD) appear to vary from left to right, albeit less dramatically compared to SHG intensity and pixel density.

### Image subdivision and texture analysis

Important morphological features of tissue can be investigated by considering the spatial relations of neighboring pixels across the computed polarimetric parameters of the region of interest. Texture analysis allows for quantification of neighboring pixel distributions, resulting in a numeric score for each of the computed texture parameters, such as contrast, correlation, entropy, angular second moment (ASM), and inverse difference moment (IDM), over the entire imaged region [29]. The imaged large area of lung tissue, in this study, spanned several millimeters in size, containing over 50 million pixels. Since the application of texture analysis over the entire image would result in a single numeric score for each of the texture parameters, it would completely disregard the subtle variations in polarimetric parameters of the imaged collagen fibers. To reveal the local variations of the polarimetric parameters, each widefield P-SHG image was subdivided into 256 smaller sub-images, each consisting of 128 × 128 pixels and covering approximately 42*μm* × 42*μm* region of the tissue.

Following the subdivision, sub-images which contained predominantly adipose tissue or blood vessels, as indicated by pathologist, were omitted from the analysis. Texture analysis was performed on each of the remaining sub-images, generating maps of texture parameters over the entire imaged region. In addition, pixel density, as well as, mean and mean absolute deviation (MAD) of the polarimetric parameter of the pixels within each sub-image were computed, resulting in 36 polarimetric and texture parameters maps that were further analyzed using unsupervised machine learning to decipher ECM modifications across various length scales from the tumor.

### Principal component analysis of parameters

In order to perform meaningful unsupervised clustering, the underlying behavior of the polarimetric and texture parameters was investigated using Principal Components Analysis (PCA). As shown in Fig. 2a, the first 4 principal components (PCs) possess the most useful information, as indicated by the dashed purple line corresponding to covariance matrix eigenvalue of 1. The first 4 PCs also explain 74.5% of the variance in the data (fig. 2b), while the first 2 PCs explain 42% and 19% of the total data variance, respectively. Therefore, we concluded that only the first 4 PCs may be utilized in unsupervised machine learning. Biplots of PC2 vs PC1 and PC4 vs PC3 allows for visualization of the extent of contribution of polarimetric and texture parameters in the 4 most important PCs, as shown in Figs. 2c-d. In biplots, the relative proportion of parameters in a given PC can be visualized with projection of the length (loading) of the eigenvectors onto the corresponding PC axis. As such, it was evident that largest contributors to PC1, and thus over 40% of the variance in the data, was mostly associated with SHG intensity and pixel density. In addition, R-Ratio and DCP correlation and IDM played an important role in separating the data. As a result, we considered PC1 to mostly represented morphological and molecular variations in the tissue. The most important factors of PC2 were SHG-CD and SHG-LD contrast and IDM, and entropy of most polarimetric parameters, which provided ample information regarding heterogeneity and disorder of collagen fiber orientation and organization. Moreover, it was clear that both and PC3 and PC4 correspond mostly to SHG intensity contrast, mean, and MAD, as well as SHG-CD and SHG-LD mean. Insets of the dashed blue bordered sections are provided for each biplot in Fig. 2e-f, showcasing the eigenvectors of the parameters with less significant contribution to the top 4 PCs. On the whole, we conclude that pixel density and SHG intensity with its associated texture parameters result in the most apparent variance in the data. As the tissue undergoes morphological changes due to the presence of tumor, degradation of collagen, or alternatively, the initial stages of fibrosis result in the appearance of collagen fragments which contribute less efficiently to the produced SHG signal. In addition, collagen fiber heterogeneity and disorder seem to cause significant variation in the data, as indicated by SHG-CD and SHG-LD contrast, IDM, entropy, as well as, DCP and R-Ratio entropy. Due to large area of the imaged tissue and the complexity of collagen organization, such variations are to be expected. As such, k-means clustering is further utilized, as described in the following section, to identify the spatial significance of the variations across the region of interest.

**Figure 2:**
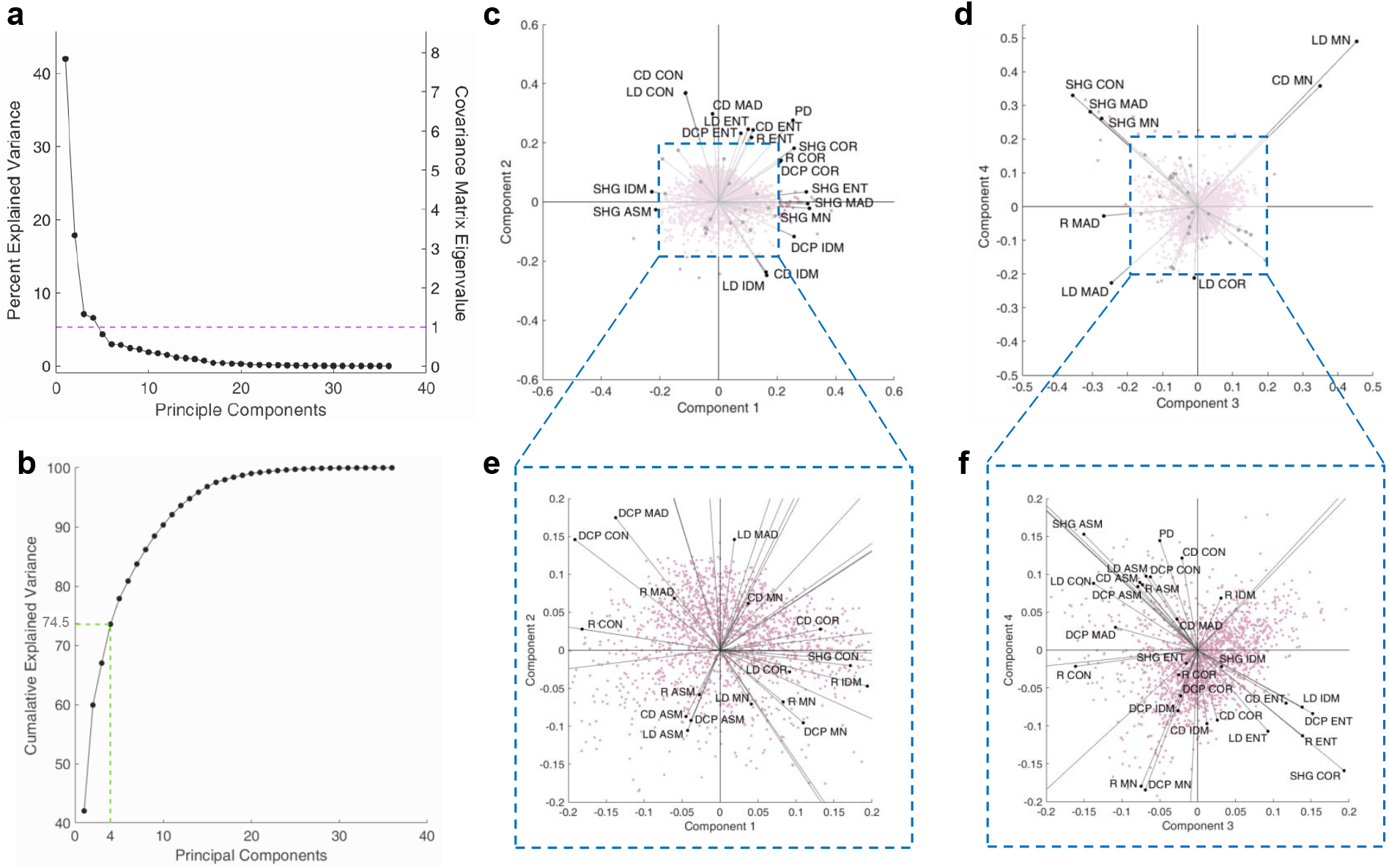
Principal Components Analysis. Percent explained variance depicts a clear elbow at PC4, also corresponding to covariance matrix eigenvalue of 1 (a). The cumulative explained variance shows that first 4 PCs account for 74.5% of the variance in the data (b). Biplots of PC2 vs PC1 (c) and PC4 vs PC3 (d), and their insets (e and f, respectively), show the most significant contributing polarimetric and texture parameters in each PC. Acronyms: PD: Pixel Density; SHG: SHG intensity; R: R-Ratio; DCP: degree of circular polarization; CD: SHG-CD; LD: SHG-LD; CON: contrast; COR: correlation; ENT: entropy; ASM: angular second moment; IDM: inverse difference moment; MN: mean; MAD: mean absolute deviation.

### Large-area tissue characterization with k-means clustering

Binary k-means clustering (k=2) was utilized for efficient consideration of the polarimetric and texture parameters in unsupervised characterization of the imaged lung tissue. As shown in Fig. 3a, k-means yielded a binary clustered map of the tissue. It was evident from the binary map that normal portion of the tissue seemed to correspond well with cluster 1, while the tumor side seemed to contain more sub-images associated with cluster 2. To better evaluate the performance of the k-means clustering results, silhouette scores of all sub-images were computed. Silhouette score measures the clustering performance by comparing the distance between each point to all points in the same cluster, with those of the opposite cluster, resulting in a numeric score ranging from -1 (completely incorrect designation) to 1 (perfect designation) (citation needed). As illustrated in Fig. 3b, k-means silhouette scores were predominantly positive, indicating adequate performance across most sub-images.

**Figure 3:**
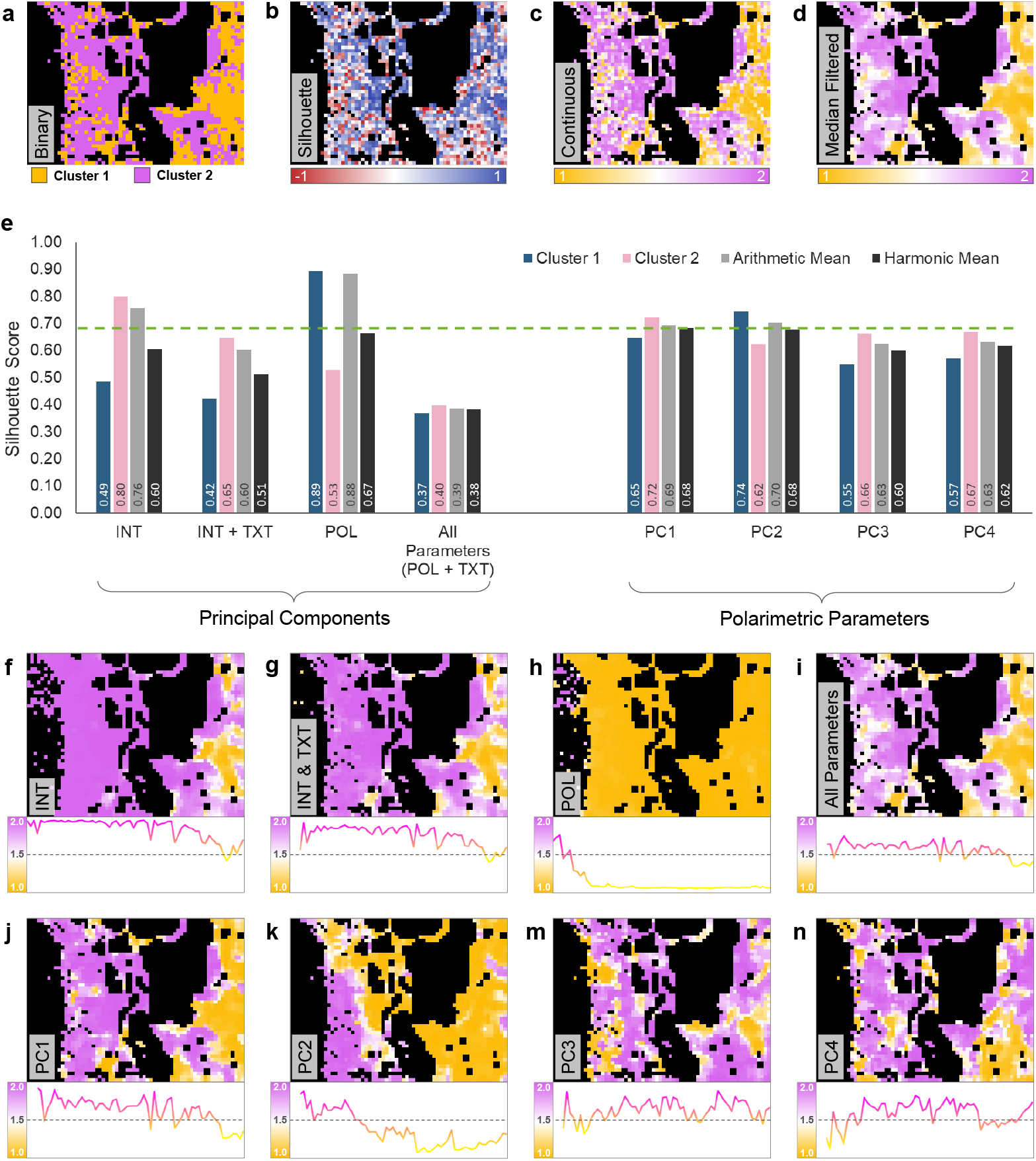
K-means clustering of lung tissue. K-means clustering was performed using all polarimetric and texture parameters for all sub-images, resulting in a binary cluster map (a). A silhouette score map of the k-means results was generated to assess clustering performance (b). The binary and silhouette maps were combined into a single map of a continuous variable ranging from 1 (cluster 1) to 2 (cluster 2) using a mathematical transformation (c). Median filter of the continuous map provides for better visualization of the clusters (d). Performance of 8 different data subsets was assessed using silhouette scores of each cluster (e). Median filtered maps of the data subsets highlight the 2 dimensional distribution of clusters, and their vertically averaged plots illustrate horizontal cluster variations (f-n).

It is important to consider the level of confidence in the association of each sub-image to its cluster, however, it is challenging to visually consider both the binary k-means clustering and the silhouette score maps when determining the location of normal and tumor tissue. As such, we have introduced a mathematical transformation that combines the cluster association and the corresponding silhouette score of each sub-image to enhance the visual assessment of the correlation between the binary and silhouette maps (details can be found in Methods). Consequently, we have illustrated the information from both binary and silhouette maps on a single map of a continuous variable (Fig. 3c) ranging from 1 (perfectly associated with cluster 1) to 2 (perfectly associated with cluster 2). When considering the continuous map, it is clear that inter-cluster boundaries which are depicted in white (mid-way between the clusters at 1.5), indicated lack of clustering confidence and association to either cluster. Furthermore, a median filtered image of the continuous map is shown in Fig. 3d, to provide a more clear image of the clustered tissue. It is evident from the median filtered map that the right side (normal) of the region of interest contained sub-images that mostly belong to cluster 1. However, moving towards left (tumor), more sub-images become associated with cluster 2, while a few islands of sub-images with normal-like collagen structures remained intact.

In order to optimize the clustering performance of the tissue, we performed k-mean for various subsets of the data, including 1) SHG intensity and pixel density (INT), 2) SHG intensity, pixel density, and corresponding texture parameters (INT+TXT), 3) all polarimetric parameters only (including SHG intensity and pixel density)(POL), and 4) the entire dataset containing all polarimetric and texture parameters and pixel density (All Parameters/POL+TXT). In addition, we performed k-means with each of the computed principal components, including 5) PC1, 6) PC2, 7) PC3, and 8) PC4. The silhouette scores of individual clusters, as well as their arithmetic and harmonic mean are shown in Fig. 3e for all considered data subsets. Data subset 1 and 3, without any texture parameters, provided the largest silhouette score arithmetic mean, however, this effect was attributed to the overwhelming designation of most sub-images into a single cluster. Median filtered maps of data subsets 1 and 3 are shown in Fig. 3f and 3h, highlighting the lack of clustering sensitivity of polarimetric parameters without their corresponding textures.

Data subset 3, corresponding to POL seems to identify the tumor mass, however, the lack of SHG signal in the region render such clustering performance potentially unreliable. When compared to data subsets 2 and 4 containing texture parameters, it is apparent that the silhouette scores of clusters were more comparable, albeit decreased overall. As such, it is important to consider the harmonic mean of silhouette scores, in place of the arithmetic mean when considering the overall clustering performance.

The k-means performed with PC1, PC2, and POL possessed the largest silhouette score harmonic mean, as indicated by the dashed green line in Fig. 3e. It is important to mention that silhouette scores were consistent for both clusters when performing k-means on PCs and gradually decreased when more PCs were included in the data subset. However. the spatial pattern of the designated clusters generally remained constant. The corresponding median filtered maps of all considered data subsets are shown in Figs. 3f-n. To better understand the cluster variations across the tissue from normal to tumor, sub-images in each vertical column of the maps were averaged. The resulting vertically averaged plots are shown below each of the maps in Figs. 3f-n. It is evident that most data subsets averaged plots are predominantly associated with one cluster as can be seen in Figs. 3f-h, or oscillate around the mid-way boundary (Figs. 3i,3m,3n), indicating lack of separation between clusters towards the normal and tumor sides. Figs. 3j-k demonstrated the most distinct variation in the horizontal direction which we shall focus on in the next section of this article.

### Tumor margin identification

Identification of the exact position of the tumor margin is crucial in characterization of the tumor and subsequent plans and procedures pertaining to excision and treatment. In this section we evaluate the utility of k-means clustering and PCA analysis on widefield P-SHG images in highlighting the tumor margin. The region of interest was investigated by the expert pathologist, E.Z., and the corresponding ground truth map of the tissue was first created. As shown in Fig. 4a, the ground truth map highlights the tumor mass and its associated margin containing individual tumor cells in red, separated by a boundary (shown in black) from the predominantly normal tissue to the right, shown in blue. All omitted regions corresponding to adipose tissue and blood vessels are transparent in the map, thus only the underlying H&E image is visible.

**Figure 4:**
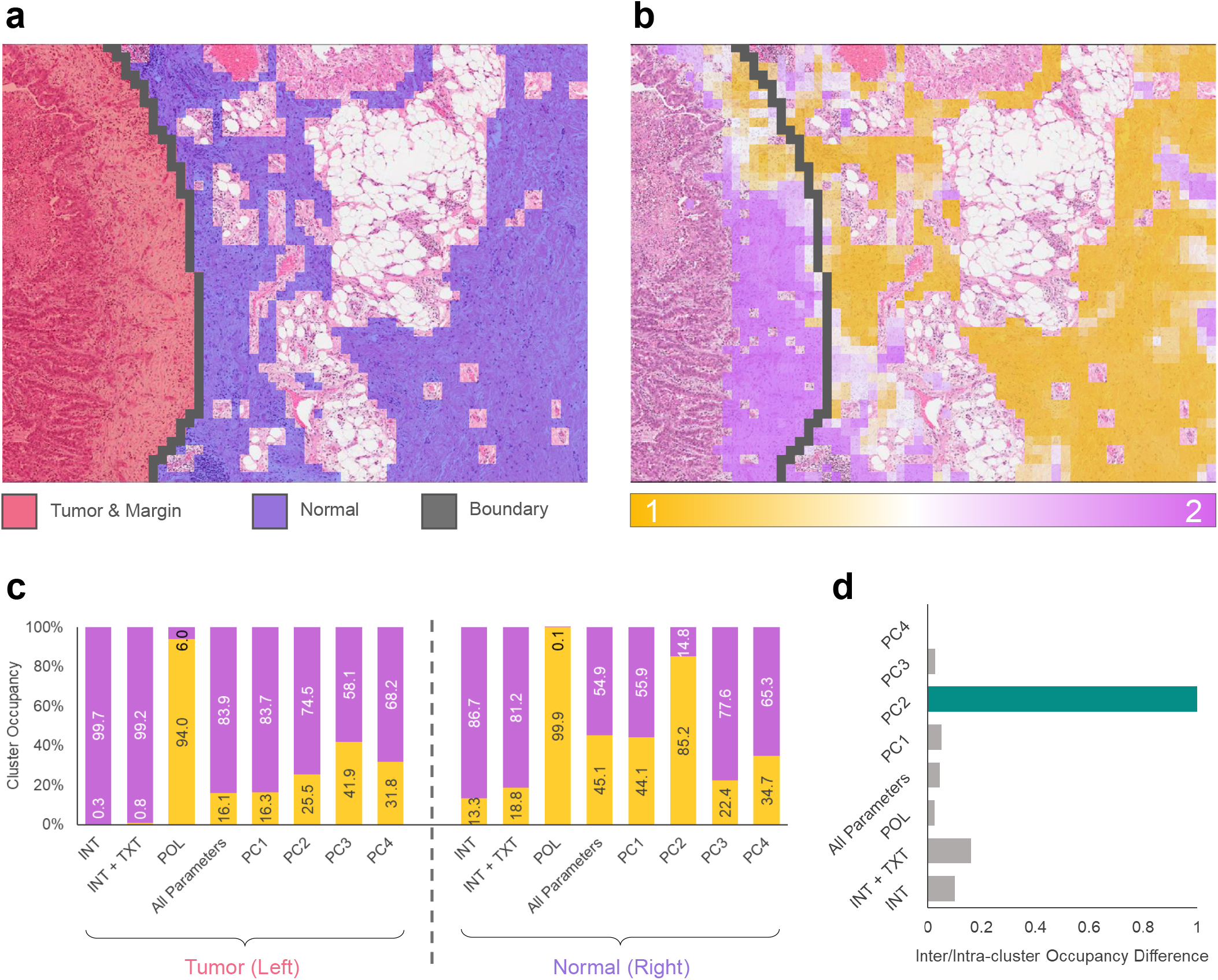
Tumor margin estimation. A ground truth map was generated to highlight the tumor mass and margin (red), as well as the normal tissue (blue) and the boundary in between (black) (a). The tumor-normal boundary was superimposed on the H&E and the k-means clustered image with PC2 data subset. The normal side mainly corresponds to cluster 1 and the tumor side is main associated with cluster 2 (b). Cluster occupancy barplot shows subset PC2 is the only data subset whose occupancy is reversed across the boundary (c). Difference between intra-cluster and inter-cluster occupancy across the boundary shows PC2 is best equipped to identify the tumor margin (d).

Using the ground truth boundary, the image was divided into tumor and normal regions. The boundary was then superimposed on each of the data subsets introduced previously. Figure 4b illustrates the superposition of the boundary, the PC2 k-means map, and the H&E image. As evident from Fig. 4b, the tumor side of the boundary mostly contains magenta-colored cluster 2 sub-images, while the normal side predominantly contains yellow cluster 1 sub-images. This behaviour was previously hinted by the crude vertically averaged plot of Fig. 3k.

To further quantify this observation, the spatial occupancy of each cluster was computed for the tumor and normal sections. The cluster occupancy was computed as a ratio of the continuous score weighted average of sub-images in each cluster, over all sub-images on each side (details can be found in Methods). As shown in Fig. 4c, the cluster occupancy of the k-means maps for each data subset were computed for both sides of the boundary. There are 2 criteria for effective differentiation of the tissue type across the boundary: 1) occupancy of one cluster should be maximized on either side of the boundary, and 2) the cluster with the larger occupancy on one side of the boundary should have a minimal occupancy on the opposite side. It is apparent that for most data subsets, the first condition is fulfilled since cluster 2 (magenta) seems to be prevalent for all but the 3rd data subset (POL), where cluster 1 accounted for the overwhelming majority of the occupancy. However, this condition is apparent on both sides of the boundary for all but one data subset, corresponding to k-means performed with PC2. As such, PC2 was deemed to be suitable in identifying the tumor boundary.

Both required criteria were combined into a single parameter, namely, the inter/intra-cluster occupancy difference (IIOD), which accounts for the inter-cluster occupancy difference on each side of the boundary, as well as, intra-cluster occupancy difference across the boundary (details can be found in Methods). The results of the IIOD computation is illustrated in Fig. 4d, where all quantities are normalized with the largest value. As determined by IIOD, it is clearly apparent that k-means performed with PC2 best identifies the boundary between the normal and tumor tissue region. As such, we conclude that linear combination of SHG-CD and SHG-LD contrast, IDM, entropy, and, DCP and R-Ratio entropy are the primary parameters in identification of the tumor margin.

## Discussion

The information provided by the structure of ECM around various pathologies such as cancer has been documented. However, application of such knowledge has not received widespread use in cancer diagnostics due to complicated scanning microscopy systems and time consuming imaging procedures. We have presented an application of the recently introduced widefield P-SHG microscopy [27] to map the spatial variation of the ECM across a large region of lung tissue in a timely manner. In this work, imaging and analysis of the entire region of interest was performed in under one hour (3 × 4 tiled image grid, where 16 polarization state images of each region was acquired in 4 minutes), enabling visualization and quantification of collagen ultrastructure across the entire imaged region. PCA provided highly optimized linear combinations of the polarimetric and texture parameters that explained the largest variance in the data, which were important for tissue type differentiation. The resulting polarimetric and texture parameters and PCs were utilized in k-means clustering to generate maps of normal- and tumor-like tissue, which were compared to the ground truth map of the tissue which highlighted the tumor mass and margin. While each parameter clustered the region differently, PC2 was found to identify the tumor margin using only the ECM features and in the absence of cellular information. As such, ECM collagen properties may be utilized as a biomarker for computer-assisted cancer diagnostics.

PC1 accounted for more than 40% of the variance in the data. Pixel density, SHG intensity and its associated texture were amongst the most significant contributors to PC1. k-means clustering performed on PC1 revealed a map similar to one attained using all parameters, albeit with much improved silhouette score. The map seemed to disregard the tumor margin by placing a prominent cluster boundary much further to the right of the tumor margin. In addition, smaller “islands” of the opposite cluster appeared throughout the image, as evident in Figs. 3i-j. It is clear that the magenta-colored cluster 2 has infiltrated the bottom right corner of the imaged region, far within cluster 1, while a few smaller cluster 1 islands areas were within the predominantly cluster 2 region to the left. Due to significance of SHG intensity in PC1, this behavior is primarily due to the strength of the SHG signal produced by the ECM. We speculate that PC1 places a greater emphasis on the weak signal produced at the center of the image, likely due to degradation of collagen fibers, or alternatively as a result of fibrosis. As such, the center of the imaged region is combined with the tumor-like tissue to form a single cluster. If so, the resulting irregular pattern could highlight the large extent of the ECM modification as a consequence of the tumor, and the appearance of potential metastatic invasion routes from the tumor site towards the normal tissue. However, further investigation is necessary for a more definitive explanation.

It is important to note that while decreased SHG intensity and pixel density were amongst the most important indicators of altered ECM collagen due to tumor development, other factors have significant impact on the produced SHG signal. Adipose tissue, blood vessels, and other sparsely populated collagenous regions in the tissue also result in decreased SHG intensity and pixel density. Fortunately, most of such regions result in a lower SHG signal than areas surrounding tumor, therefore, fall below the minimum SNR level of 1 and can be discarded from the analysis. There were a number of blood vessels and significant areas of adipose tissue throughout the region of interest. In order to eliminate the potential inclusion of such areas in the analysis, all sub-images that coincided with adipose tissue and blood vessels was discarded.

In future developments, the previously introduced chiral second order nonlinear optical susceptibility ratio, C-Ratio 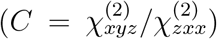, may be computed and used in tissue clustering. C-Ratio has been shown in scanning P-SHG microscopy to probe the chiral nature of collagen fibers, while providing important information on collagen polarity and 3-dimensional organization [9, 43]. In addition, H&E images may be used to introduce stain color variations to the analysis and enable automated rejection of sub-images with adipose tissue. Moreover, third-harmonic generation (THG) images can also be introduced to allow for inclusion of cell morphology in the analysis routine. Cellular information provided by the H&E and THG images complement the ECM-specific P-SHG polarimetric and texture parameters, resulting in reduced false positive and false negative rates.

## Materials and Methods

### Widefield P-SHG microscope and procedure

The custom made widefield P-SHG microscope used in this word was described previously [27]. Briefly, a high-power amplified laser (PHAROS PH1-15W, Light Conversion) with 1.3 W power, 1030 nm wavelength, 100 kHz repetition rate, and 13*μ*J of energy per pulse was coupled to the microscope to illuminate a large area of the tissue. The laser beam was first passed through an infrared polarizing beam splitter (PBS102, Thorlabs), followed by an infrared liquid crystal variable retarder (LCVR) (LCC1223-B, Thorlabs), to attain its initial polarization state. A 30-mm achromatic doublet (AC254-030-AB, Thorlabs) was used to focus the beam just beneath the sample, to allow for adjustment of the illumination area through vertical translation. The produced SHG signal and the fundamental beam was collected by a Plan-Neofluar 20×*/*0.50 air objective (420350-9900-000, Zeiss). The collected light then passed through a visible LCVR (LCC1223-A, Thorlabs) and a visible polarizing beam splitter (PBS251, Thorlabs) to determine the outgoing polarization state of SHG. The SHG signal was separated from the fundamental beam using a dichroic mirror (FF662-FDi02-t3-25×36, Semrock). The separated SHG signal was filtered by two BG40 colored glass filters (FGB37-A, Thorlabs) and a 515/10nm interference filter (65-153, Edmund Optics). The filtered light was projected onto a sCMOS camera (ORCA-Flash 4.0 V2, Hamamatsu).

A total of 16 polarization state combinations with 4 incoming light polarizations and their corresponding 4 outgoing SHG signal polarizations (left circular polarized (LCP), right circularly-polarized (RCP), vertically linearly polarized (VLP), and horizontally linearly polarized (HLP)) were used to carry out P-SHG microscopy imaging. The P-SHG maps of the region of interest were constructed by tiling 12 images (3 × 4 grid), which were acquired in 48 minutes. Each image was acquires in 4 minutes, with 10 seconds of camera exposure and 10 seconds of polarization switching.

Once the images were acquired, Stokes Mueller Polarimetry was employed to compute 3 Stokes vector elements for each of the 4 incoming polarization states (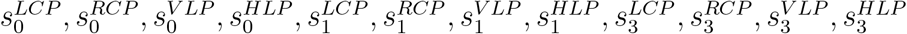, where the superscripts indicate the incoming polarization state and *s*_0_, *s*_1_, and *s*_3_ represent the normalized total SHG intensity, normalized difference in linearly polarized SHG intensity, and normalized difference in circular polarized SHG intensity, respectively). The computed Stokes vector elements were combined to create 5 polarimetric parameters including the average circularly-polarized SHG intensity expressed as

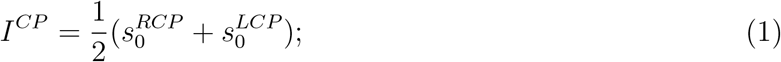

a ratio of two achiral second order nonlinear optical susceptibility tensor elements (R-Ratio) [50]

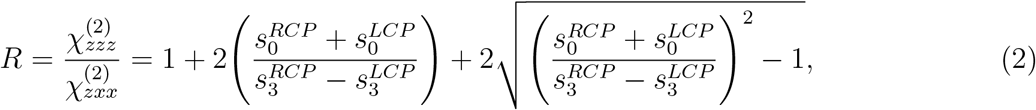

where z points along the fiber axis; degree of circular polarization (DCP) [51]

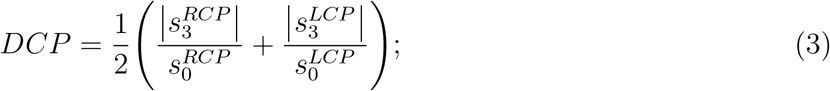

as well as SHG circular and linear dichroism (SHG-CD and SHG-LD, respectively), expressed [27, 48, 49, 50]

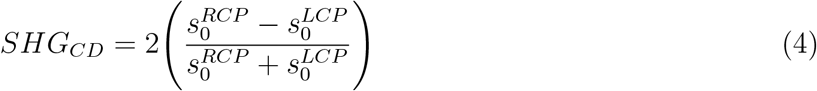

and

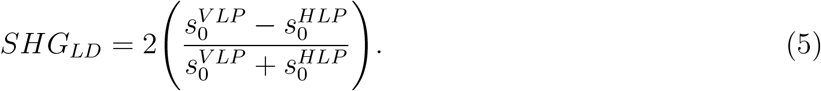

### Texture analysis

Texture parameters were extracted from the grey-level co-occurrence matrix (GLCM), for a given region of interest [29]. The (*i, j*)^*th*^ element of the GLCM indicated the number of occurrences of a pixel at grey level *i* followed by a pixel at grey level *j*. For nearest neighbour analysis (distance factor *d* = 1), a separate GLCM was created for 4 angles corresponding to *θ* = {0°, 45°, 90°, 135°}. These GLCMs were then averaged to create an overall direction-independent GLCM. The resulting GLCM was then normalized to obtain the probability density function of finding grey-level pairs, *P_d,θ_*(*i, j*), to calculate the 5 texture parameters of interest (as described by Haralick et. al [29]). The texture parameters of interest were contrast, correlation, entropy, angular second moment (ASM) and inverse difference moment (IDM).

Contrast, *ν*, is a measure of grey-level variation between pixel pairs. It can be used to probe collagen fiber density in the tissue:

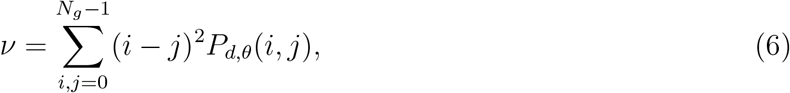

where *N_g_* is the number of gray levels, representing the discretization level of continuous polarimetric parameters.

Correlation, *ρ*, quantifies a linear dependence between grey-level pairs. It is given by:

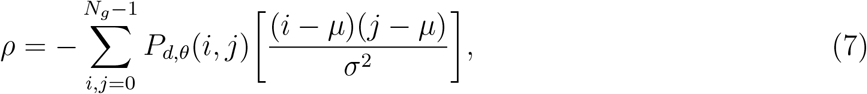

where *μ* is the mean and *σ* is the standard deviation of the grey levels. For perfectly negatively and perfectly positively correlated images, the correlation would be -1 and 1, respectively. In the context of widefield P-SHG microscopy, correlation would be indicative of the presence of well-defined structures and patterns in the images.

Entropy, *S*, given by:

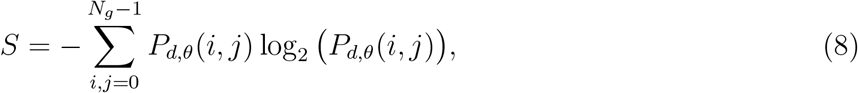

is a measure of the level of disorder - or lack of spatial organization - of the grey levels. When disorder increases, entropy also increases. However, entropy is also indicative of the size of pixel clusters, which, in the context of widefield P-SHG, often represent groupings of collagen fibers. Conversely, angular second moment (ASM) is a measure of orderliness. ASM ranges from 0 to 1, where 1 is indicative of a completely uniform image. ASM also varies with the size of pixel clusters of comparable values and is a measure of the average size of similar collagen fiber bundles in an image. ASM is expressed as:

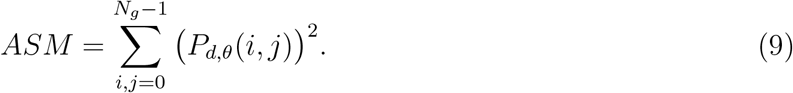

Finally, inverse difference moment (IDM) describes the homogeneity of an image. Similar to ASM, the range of IDM is from 0 to 1, where 1 is characteristic of a completely uniform image. IDM is provided by:

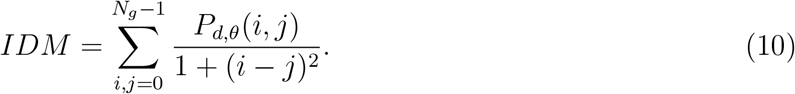

The size of the GLCM is dependent on *N_g_*, therefore, a custom program was written in MATLAB to perform texture analysis as described previously [27]. Often, widefield P-SHG images contain dark regions between collagen fibers, which result in significant background noise and lead to highly skewed texture distributions and reduced differentiation power. To accommodate, the background pixels were removed from analysis, allowing for texture parameters to reflect the structure of signal-producing entities, such as collagen-fibres, and disregard irrelevant data.

### PCA and k-means clustering application

Each of the acquired P-SHG images were subdivided into 256 sub-images, in order to create a map of polarimetric and texture parameters throughout the region of interest. While treating each sub-image as a separate data point, the resulting dataset was comprised of 3072 points, 2381 of which possessed a large enough SHG signal (SNR *>* 1) and were used in further analysis using unsupervised machine learning.

### PCA

Principal component analysis was employed to identify polarimetric and texture parameters which corresponded to the largest variation in the data. Prior to the application of PCA, the dataset was standardized, such that mean of each parameter was subtracted and subsequently divided by their respective standard deviations. Consequently, only true variations between data points of each parameter accounted for the variance of the data, and differences in scale and units were omitted. PCA was performed on the standardized data, using MATLAB, and the resulting principal components were visualized using biplots. Moreover, figures of explained variance and covariance matrix eigenvalues as a function of principal components were used to identify the most significant principal components.

### Combination of binary k-means clustering with silhouette score

Eight different subsets of the data were further used in binary k-means clustering (*k* = 2) with MATLAB. The results were rearranged to spatially construct the binary clustered map of the tissue. Silhouette score of each clustered sub-image was computed to evaluate the clustering performance. The silhouette score compares the distance of a single data point with points in the same and opposite cluster, resulting in a numeric value that ranges from -1 for completely incorrect cluster association, to 1 for perfectly correct cluster association. A map of the silhouette score was generated to display clustering performance.

It is challenging to simultaneously visualize the binary clustered map and the silhouette map. As such, the following mathematical transformation is introduced to combine both binary cluster association (*B*), and the silhouette scores (*S*) of each point into a continuous cluster association (*C*):

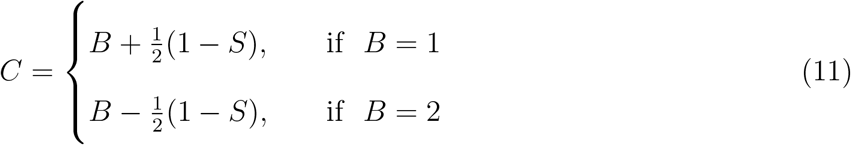

The continuous parameter, *C*, ranges from 1 to 2, corresponding to points perfectly associated with clusters 1 and 2, respectively. The range between 1 and 2 accounts for non-perfect silhouette scores, and the inter-cluster boundary lies at 1.5, whose corresponding point do not belong in either cluster. It is important to mention that this transformation also accounts for points with negative silhouette scores, by moving such points to the opposite cluster.

The resulting continuous variable maps often contained rough spatial variations. Median filters of the maps were computed to generate smoothly varying approximate clustered images for better visualization. The median filter computation was performed in MATLAB, where the value of each pixel was replaced by the median of itself and its 8 nearest neighbors. Dark sub-images corresponding to rejected data points were left as NaNs.

### Determination of inter/intra-cluster occupancy difference

To determine the performance of the algorithm in identification of the tumor margin, the occupancy of the 2 clusters is computed on each side of the boundary. In the simplest case with perfect clustering silhouette score of 1, the cluster occupancy of cluster 1 reflects the number of sub-images belonging to cluster 1 divided by in the region of interest, while the same would be true for cluster 2. However, when considering non-ideal clustering performance, a weighted average of the continuous parameter, *C*, is required:

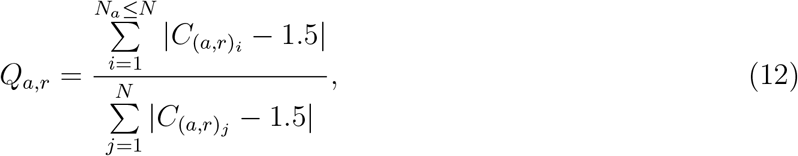

where *Q_a,r_* denotes the cluster occupancy of cluster *a, C*_(*a,r*)_ represents the continuous parameters of cluster *a, N* is the total number of sub-images, and *N_a_* is the number of sub-images of cluster *a* in the region *r*. Note that the continuous variable is shifted by 1.5 in order to set the range symmetrically across 0, from -0.5 to 0.5. Subsequently, the absolute value of the shifted *C* is computed and used as weight in the above equation. This procedure correctly associates sub-images with negative silhouette scores with the opposite cluster.

The inter/intra-cluster occupancy difference (IIOD) is introduced to evaluate the similarity of each cluster with any given area of the imaged region. The IIOD accounts for the difference between the occupancy of clusters within each area (inter-cluster), as well as, the difference between the occupancy of the same cluster in different areas in the region of interest (intra-cluster). Therefore, for a binary system (2 clusters) in 2 different areas (e.g. tumor on the left and normal on the right, separated by a boundary) IIOD can be expressed as:

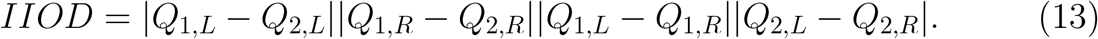

According to the above equation, the maximum IIOD score of 1 is attained in the case where each cluster is only found on either the left or right of the boundary. Conversely, IIOD will reach a minimum of 0 when clusters exhibit 50% occupancy or when only one cluster accounts for the entire imaged region. Thus, IIOD of k-means clustering performed with various data subsets may be compared to highlight the most important polarimetric and texture parameters, as the corresponding morphological features, that identify the normal and tumor sections of the imaged region.

### Sample preparation

The tissue sample was first fixed in paraffin, then cut into 5 *μ*m thick sections, and mounted on glass slides. The tissue sections were stained with hematoxylin and eosin and imaged with a bright-field microscope scanner (Aperio Whole Slide Scanner, Leica Biosystems) for histopathology investigations. A 2mm × 2.7mm tissue region encompassing tumor (left side of the region) and normal-like tissue (right side of the region) was selected by expert pathologists (E.Z. and M.T.) to investigate ECM variations in the tumor margin.

### Ethics Statement

The sample of human normal and primary non-small cell lung carcinoma tissues were obtained with written informed patient consent, and the study was approved by the Research Ethics Board of University Health Network, Toronto, Canada, in accordance with relevant guidelines and regulations and in compliance with the tenets of the Declaration of Helsinki.

## Data availability

All raw and processed image files analysed during this study are available from V.B. upon request.

## Acknowledgements

This work was supported by Natural Sciences and Engineering Research Council of Canada (NSERC) (RGPIN-2017-06923, DGDND-2017-00099, CHRPJ 462842-14), the Canadian Institutes of Health Research (CIHR) (CPG-134752), and European Regional Development Fund with the Research Council of Lithuania (01.2.2.-LMT-K-718-02-0016). We thank Light Conversion for providing the laser for our experiments.

## Author Contributions

V.B., B.C.W., M.T, K.Y. and K.M. designed the experiment. L.K. optimized the laser source. R.N, M.T prepared the H&E-stained tissue slide, M.T and E.Z selected region of interest for widefield P-SHG imaging. K.M. and L.U.C. calibrated the microscope and imaged the lung tissue sample. K.M. analyzed data, performed principal components analysis, and k-means clustering. Y.K. performed texture analysis. K.M., L.U.C., Y.K., A.G., and V.B. interpreted the results and wrote the article draft. All authors contributed to the manuscript.

